# Encapsulated allografts preclude host sensitization and promote ovarian endocrine function in ovariectomized young rhesus monkeys and sensitized mice

**DOI:** 10.1101/2021.06.10.447598

**Authors:** James R. Day, Colleen L. Flanagan, Anu David, Dennis J. Hartigan-O’Connor, Mayara Garcia de Mattos Barbosa, Michele L. Martinez, Charles Lee, Jenna Barnes, Evan Farkash, Mary Zelinski, Alice Tarantal, Marilia Cascalho, Ariella Shikanov

## Abstract

Transplantation of allogeneic donor ovarian tissue holds great potential for female cancer survivors who often experience premature ovarian insufficiency. To avoid complications associated with immune suppression and to protect transplanted ovarian allografts from immune-mediated injury, we have developed an immuno-isolating hydrogel-based capsule that supports the function of ovarian allografts without triggering an immune response. Encapsulated ovarian allografts implanted in naïve ovariectomized BALB/c mice responded to the circulating gonadotropins without direct revascularization and maintained function for 4 months, as evident by regular estrous cycles and presence of antral follicles in the retrieved grafts. Repeated implantations of encapsulated mouse ovarian allografts did not sensitize naïve BALB/c mice in contrast to non-encapsulated controls, which was confirmed with undetectable levels of allo-antibody. Further, encapsulated allografts implanted in hosts previously sensitized by implantation of non-encapsulated allografts restored estrous cycles similarly to our results in naïve recipients. Next, we tested the translational potential and efficiency of the immune-isolating capsule in a rhesus monkey model by implanting encapsulated ovarian auto- and allografts in young ovariectomized animals. The encapsulated ovarian grafts survived and restored basal levels of urinary estrone conjugate and pregnanediol 3-glucuronide during the approximate 4-5 month observation period. We demonstrate, for the first time, that encapsulation of ovarian allografts prevents sensitization and protects the allograft from rejection in young rhesus monkeys and in sensitized mice.

## Introduction

Over the past decades, childhood cancer survival rates have increased substantially reaching over 85% due to advances in new therapies. However, the same cancer therapies can potentially lead to a myriad of health complications in childhood cancer survivors ^1–4^. One of the most common complications for female cancer survivors is premature ovarian insufficiency (POI) caused by gonadotoxic treatments ^5–7^. POI leads to depletion of the follicular pool, resulting in infertility and disruption of ovarian endocrine function. POI is particularly detrimental in children with a cancer diagnosis prior to puberty where hormonal loss can impact the maintenance of homeostasis in parallel with other endocrine organs ^8–10^. Changes during puberty also promote physical and psychological development into adulthood and can determine height, bone health, insulin responsiveness, lipid metabolism, cardiovascular health, and cognition. These changes are orchestrated by the pulsatile secretion of gonadotropin releasing hormone (GnRH) from the hypothalamus, which regulates the release of gonadotropins, and luteinizing (LH) and follicle-stimulating (FSH) hormones from the pituitary. FSH and LH stimulate the production of ovarian hormones and peptides including estradiol, androstenedione, progesterone, inhibins A and B, activin, and follistatin. The production of ovarian hormones is tightly regulated by a negative feedback loop that inhibits the production of GnRH and FSH in the brain and pituitary ^11,12^, known as the hypothalamic-pituitary-gonadal (HPG) axis, which is crucial for development of reproductive, musculoskeletal, cardiovascular, and immune systems as well as endocrine regulation ^13,14^. In young patients with POI, the HPG axis is disrupted due to deficiencies in ovarian hormones, which leads to imbalances across the endocrine system resulting in comorbidities such as suboptimal bone development, metabolic changes, and abnormal fat deposition ^14–16^.

Currently, the only clinically available option to treat POI in adolescent girls to initiate puberty is hormone replacement therapy (HRT), which delivers gradually increasing, yet fixed, amounts of estrogen ^16,17^. Hormone replacement therapy was originally designed to treat postmenopausal symptoms in women and thus the impact on children is not well-established ^17,18^. Additionally, HRT delivers only a fraction of ovarian hormones in a non-pulsatile manner which does not mimic physiological puberty and the complexity of the HPG axis. In turn, non-physiologic HRT given to induce puberty in girls with POI can also lead to premature closure of growth plates, cessation of bone growth, and metabolic imbalances ^19,20^. Auto-transplantation of cryopreserved ovarian tissue banked prior to anti-cancer treatments is a potential option under investigation. Transplanted autologous ovarian tissue restored endocrine function and the transition to puberty in girls ^6,21,22^, and has led to more than 100 babies born to either identical twin sisters or cancer survivors who cryopreserved their ovaries ^23–27^. However, this option is associated with the risk of re-introducing malignant cells potentially harbored within the ovary, particularly in patients with hematological malignancies, the most common childhood cancer ^5,28,29^. Although malignant cells can be detected with some cancers, there has been no universal and safe protocol established to screen ovarian tissue contributing an uncertain risk of re-seeding in those undergoing autologous transplantation. In addition, most childhood cancer survivors do not cryopreserve tissue prior to treatment ^10,30,31^.

We hypothesized that donor allogeneic ovarian tissue, implanted subcutaneously will respond to circulating gonadotropins and secrete ovarian hormones mimicking the physiological process while mitigating the risk of re-introduction of cancer cells. Previously we developed and optimized a novel dual-layered poly(ethylene glycol)-based (PEG) capsule that sustains survival and function of allogeneic mouse ovarian tissue while preventing rejection and supporting ovarian development in immune competent mice ^32–36^. The capsule consists of a degradable hydrogel core that provides a supportive environment for the implanted tissue allowing the cyclical changes associated with folliculogenesis, surrounded by a non-degradable hydrogel shell that allows free diffusion of essential metabolites and nutrients while preventing rejection by blocking the infiltration of host immune cells. Our studies in mice have demonstrated that encapsulated ovarian allografts undergo folliculogenesis, producing ovarian hormones in a regulated physiological manner, restoring endocrine function in a mouse model of POI without sensitizing the immune system of the host ^34,37,38^.

In a clinical setting, each patient may require recurring allo-transplantations due to a finite number of follicles transplanted in each graft due to size limitations of the graft. Recurring allografts may sensitize the host and as a result accelerate rejection and shorten the duration of graft function. In this study, we first investigated whether the capsule immune-protects the ovarian allograft in a pre-sensitized murine host that otherwise rejects non-encapsulated allografts within days of transplantation ^39,40^. Second, we implanted encapsulated auto- and allografts in young ovariectomized rhesus monkeys, which are anatomically and physiologically closer to humans, towards translation of this approach ^41–46^. Our findings suggest that the auto- and allogeneic ovarian tissues implanted in ovariectomized mice and rhesus monkeys survived in the capsule, secreted estradiol and progesterone, and do not elicit immune responses and rejection.

## Results

### Immuno-isolating capsule prevents sensitization of the host: Studies in a murine model

Implantation of non-encapsulated allogeneic ovarian tissue induced an immune reaction mediated by T and B cells, resulting in sensitization of the host against the donor with production of allo-specific antibodies and cell-mediated destruction of the graft tissue (**Figure 1A – study design, 1B, C**). As expected, implantation of non-encapsulated ovarian allografts elicited an immune response when implanted subcutaneously resulting in a 23-fold increase in allo-specific IgG (**Figures 1B and C**). Allo-specific IgG antibodies gradually increased in 4 of 7 mice from undetectable post-implantation to an average of 601 mean fluorescence intensity (MFI) on day 28 and an average of 840 MFI on day 42. To block immune-mediated graft damage, we created a barrier between the allograft and the host to minimize the host exposure to allo-antigens and prevent stimulation of allo-immunity and graft damage owing to pre-existent allo-immunity. We then assessed if implantation of encapsulated allografts allowed repeat implantation without the risk of rejection (**Figure 1D – study design**). In contrast, encapsulated ovarian allografts did not sensitize recipients suggested by undetectable levels of circulating allo-antibodies. Following the first and subsequent implantation of encapsulated ovarian allografts (**Figure 1D**) the levels of circulating allo-specific IgGs remained at the pre-implant level, ranging from between non-detectable to 138 MFI for up to 60 days post-implantation (**Figures 1E and F**). Because the second implantation of encapsulated allografts did not increase circulating IgG allo-antibodies, we concluded that neither primary nor secondary implantation of encapsulated allografts sensitized recipients.

**Figure 1:**
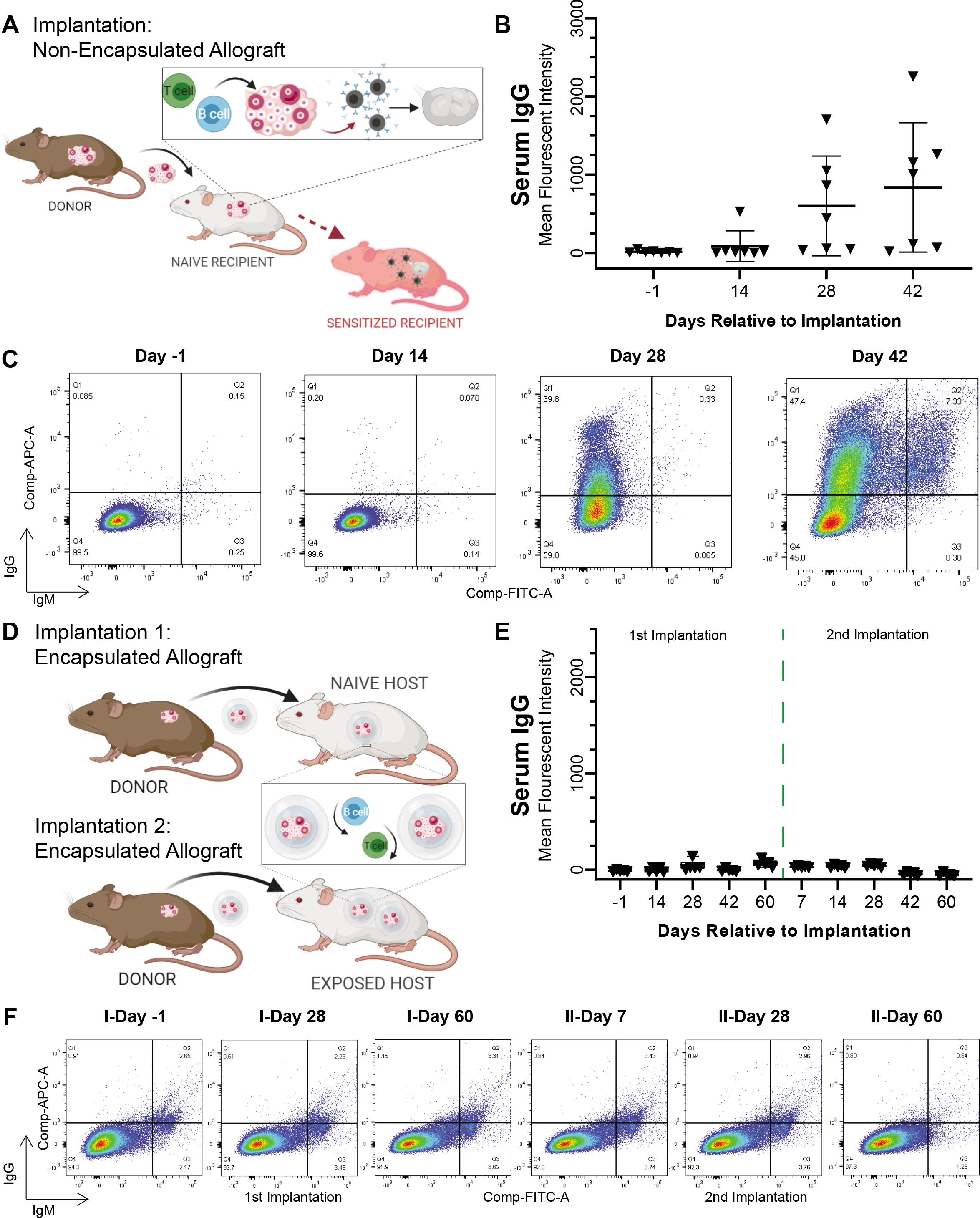
Implantation of encapsulated and non-encapsulated ovarian allografts in immune competent mice. (**A**) Schematic for implantation of non-encapsulated ovarian allograft, which causes sensitization of the recipients, production of allo-antibodies, T cell infiltration, and rejection. (**B**) Flow cytometric analysis of serum obtained from recipient mice with non-encapsulated ovarian allografts indicated average mean fluorescence intensity (MFI)±SD of allo-specific IgG over time after implantation (n=7 mice per timepoint). (**C**) Representative flow cytometry plots of binding of serum allo-specific antibodies from recipients of non-encapsulated ovarian allograft after implantation for up to 42 days. (**D**) Schematic for implantation of encapsulated ovarian allograft, which blocks interaction between the graft and recipient’s immune system and prevents sensitization of the recipient. (**E**) Average MFI±SD of allo-specific IgG over time, obtained from serum of recipient mice by flow cytometry, following implantation of two consecutive encapsulated ovarian allografts. (**F**) Representative flow cytometry plots detecting serum allo-specific antibodies from recipients implanted with two consecutive encapsulated ovarian allografts.

### Immuno-isolating capsule prevents rejection of allogeneic ovarian tissue and supports endocrine function in a murine model

Graft morphology and function were compared between mice implanted with non-encapsulated (**Figure 2A**) and encapsulated allografts (**Figures 2D-F**). The non-encapsulated ovarian allografts were resorbed based on gross examination and histology. Histological analysis of the tissue retrieved from the implantation site demonstrated necrosis, no surviving healthy follicles, and loss of distinctive ovarian morphology (**Figure 2A**). All ovariectomized mice that received non-encapsulated ovarian allografts resumed estrous cyclicity one-week post-implantation suggesting restoration of ovarian function, but cyclicity ceased after 4 weeks and remained as persistent diestrus with failed ovarian function (**Figure 2B** is a representative plot of estrous activity in an individual mouse and **Figure 2C** shows the data combined for all mice in the control group). Allograft failure was accompanied by an increase in circulating IgG allo-antibodies (**Figures 1B and C**). In contrast, all ovariectomized mice implanted with encapsulated ovarian allografts resumed cyclicity after 2 weeks which persisted for 60 days post-implantation and until the grafts were retrieved (**Figures 2G and H**). Histological analysis of the allografts revealed healthy ovarian tissue with multiple follicles at different developmental stages (**Figures 2D and E**). Although our results suggest that the encapsulated ovarian tissue did not sensitize the host, in a more stringent test we asked if a second allograft from the same source was compatible with function. To address this question, we first implanted mice with encapsulated ovarian allografts for 60 days followed by a second encapsulated ovarian allograft for an additional 60 days. Multiple healthy developing follicles up to the antral stage (**Figure 2F**) were present in the allografts retrieved from the second implant, confirming that folliculogenesis was not affected by the first implant. All mice resumed estrous activity by 2 weeks following implantation and continued cycling through the second period of 60-days post-implantation suggesting restoration of ovarian endocrine function. Because encapsulated allografts in the second round functioned as well as primary encapsulated allografts, we concluded that encapsulation may allow multiple implants without risking loss of function due to an immune response.

**Figure 2:**
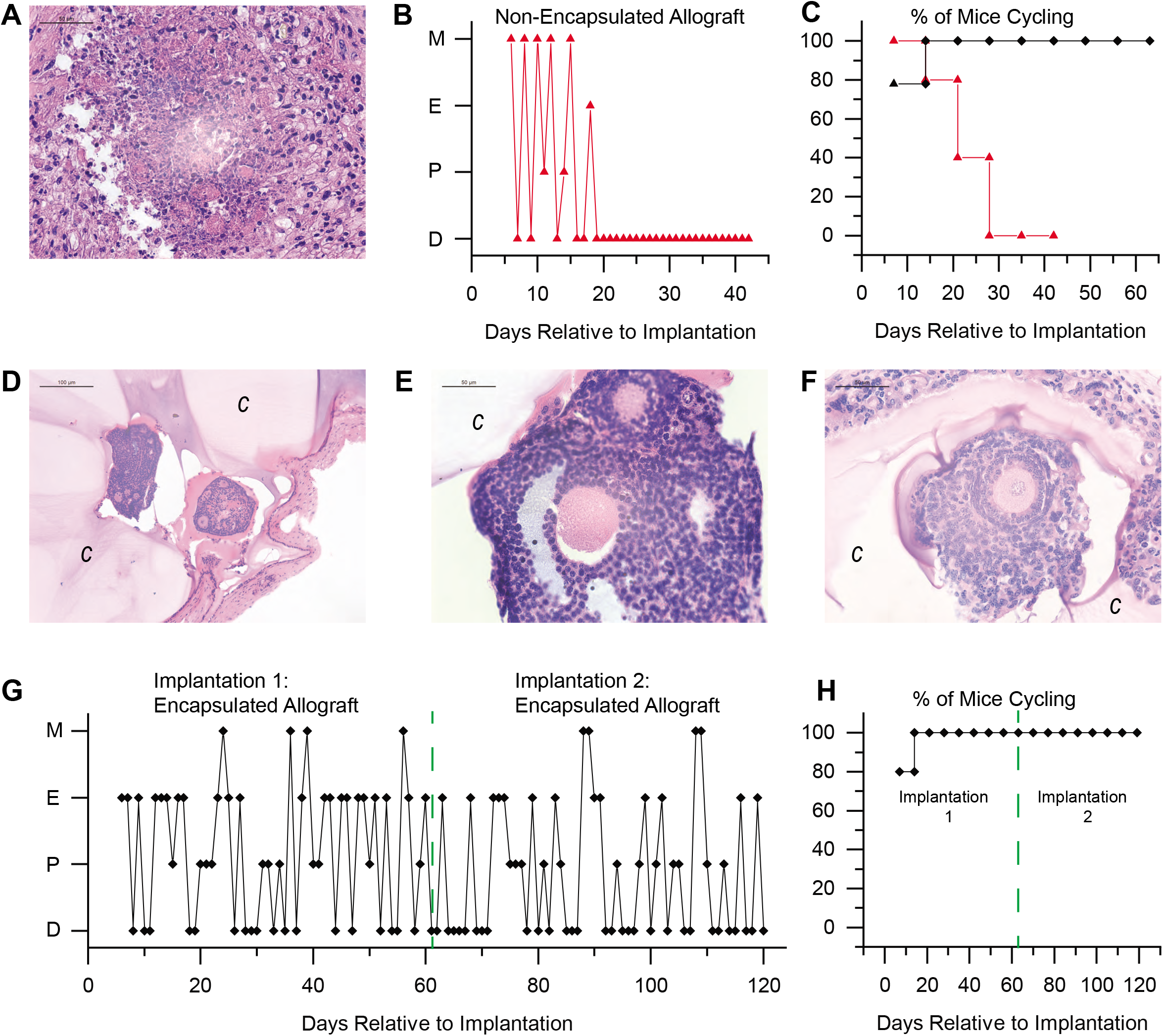
Encapsulation of ovarian allografts preserves the structure and function of ovarian tissue. (**A**) Representative histology of implanted non-encapsulated ovarian allograft. No phenotypically intact follicles were identified 6 weeks post-implantation. (**B**) Representative plot of estrous activity of mice receiving a non-encapsulated ovarian allograft; Metaestrus-M, Estrus-E, Proestrus-P, Diestrus-D. (**C**) Estrous cyclicity in mice implanted with non-encapsulated (red, n=7) and encapsulated (black, n=5) ovarian allografts. (**D**, **E**, **F**) Ovarian allografts retrieved after the first (**D**, **E**) and second round (**F**) of implants were completely surrounded by the hydrogel capsule and isolated from the host. Multiple follicles at various developmental stages (primordial through antral) were present. (**G**) Representative plot of estrous activity in mice receiving two rounds of encapsulated ovarian allografts. (**H**) Continued estrous cyclicity following two rounds of encapsulated ovarian allografts (n=5).

### Encapsulated allografts restored ovarian endocrine function and were shielded from rejection in sensitized murine hosts

We demonstrated that encapsulated ovarian allografts implanted in naïve mice were protected from immune rejection even after multiple implantations. However, the question remained whether the encapsulated ovarian allografts were protected in a host that had been previously sensitized (**Figure 3A**). To answer this question, we first implanted a non-encapsulated ovarian allograft, which sensitized the host and caused production of allo-antibodies. The sensitization was confirmed by high levels of circulating alloantibodies (on average >800 MFI, **Figures 1B, 3F**) and by cessation of estrous cycles approximately 3 weeks post-implantation (**Figures 3D and E**). We then implanted encapsulated ovarian allografts in the sensitized hosts. All sensitized recipients of encapsulated allografts resumed estrous cyclicity by 2 weeks post-implantation and continued cycling throughout the entire implantation period (**Figures 3D and E**). In spite of the presence of detectable circulating allo-antibodies persisting from prior implantation of non-encapsulated ovarian tissue, histological analysis of the encapsulated ovarian allografts 60 days post-implantation revealed fully encapsulated ovarian tissue with multiple healthy developed follicles at preantral and antral stages similar to observations in encapsulated ovarian tissue implanted in naïve mice (**Figures 3B and C**).

**Figure 3:**
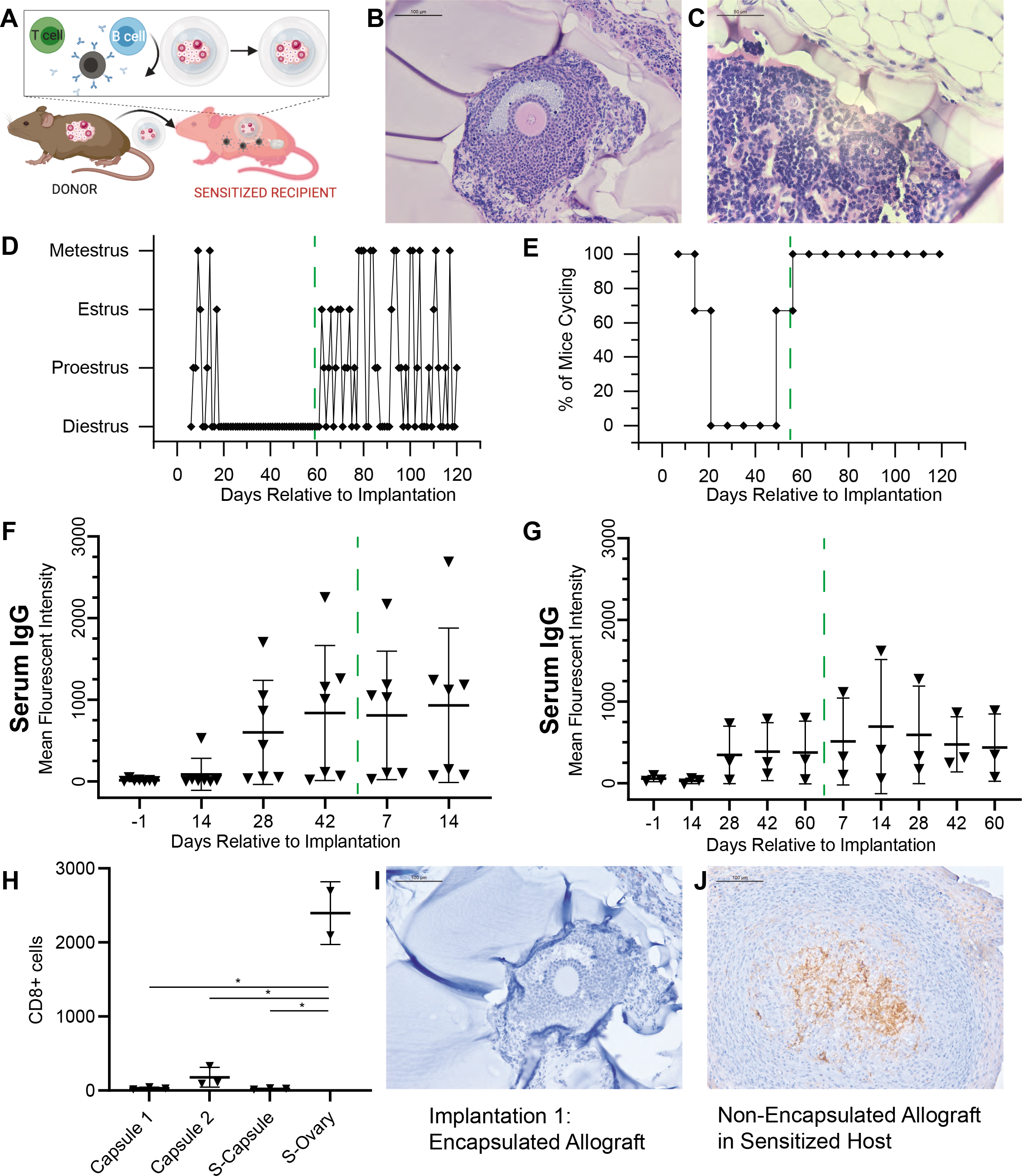
Encapsulated allografts restored ovarian endocrine function and were protected from rejection in sensitized hosts. (**A**) Schematic for implantation of non-encapsulated ovarian allograft with production of allo-antibodies, T cell infiltration, and rejection, followed by implantation of an encapsulated allograft. (**B**, **C**) Representative histology of encapsulated ovarian allografts retrieved after implantation demonstrating that the tissues were surrounded by the hydrogel capsule and isolated from the host. Multiple follicles at various developmental stages, antral (**B**) and preantral (**C**) were present. (**D**) Representative plot of estrous activity of mice receiving a non-encapsulated followed by an encapsulated ovarian allograft. (**E**) Summarized estrous cyclicity for all the mice implanted with non-encapsulated followed by encapsulated ovarian allografts (n=3). (**F**) Average MFI±SD of allo-specific IgG in mice after implantation of two consecutive non-encapsulated ovarian allografts (n=7). (**G**) Average MFI±SD of allo-specific IgG in mice after implantation of a non-encapsulated allograft followed with an encapsulated allograft after 42 days (n=3). (**H**) CD8+ T cells, identified via IHC, were present in the retrieved encapsulated allografts after first (Capsule 1) and second implantations (Capsule 2), in a retrieved encapsulated allograft implanted in a sensitized mouse (S-Capsule), and in a retrieved non-encapsulated allograft implanted in a sensitized host (S-Ovary). (**I**) Representative anti-CD8 IHC staining of encapsulated allografts for all encapsulated groups, (**J**). Representative anti-CD8 IHC staining of the non-encapsulated allograft.

Naive animals implanted with non-encapsulated ovarian allografts maintained elevated circulating allo-specific IgG, on average, 600 MFI at 28-days post-implantation (**Figures 3F and G**). A secondary follow-up implant with non-encapsulated ovarian tissue in sensitized recipients increased allo-specific IgG antibodies 7 days after implantation surpassing 810 MFI, suggesting a memory response (**Figure 3F**). Immunohistochemical analysis of the non-encapsulated allografts (S-Ovary) demonstrated significant infiltration of CD8^+^ T cells, consistent with rejection (**Figures 3H and J**).

In contrast, following the first implantation of encapsulated allograft (Capsule 1), the second implantation of the encapsulated allograft (Capsule 2), and implantation of an encapsulated allograft following sensitization (S-Capsule), the allografts had no CD8^+^ cell infiltrate inside the graft (**Figures 3H and I**). Figure 3I is a representative histological image of the immunohistochemical (IHC) staining for CD8 that was typical of all encapsulated allografts in these groups, illustrating the lack of CD8^+^ T cells within the allograft. Importantly, following the secondary implantation of an encapsulated allograft in a sensitized host (S-Capsule), the encapsulated allografts had no CD8^+^ T lymphocyte infiltration (**Figures 3H and I**), in spite of high levels of circulating allo-specific IgG **(Figure 3G**). These observations suggest that encapsulated ovarian allografts were not rejected despite immune pre-sensitization.

### Encapsulation and implantation of nonhuman primate ovarian tissue in ovariectomized young rhesus monkeys

We performed two rounds of subcutaneous implantations of encapsulated autologous and allogeneic ovarian tissue fragments in ovariectomized young rhesus monkeys (**Figure 4**).

**Figure 4:**
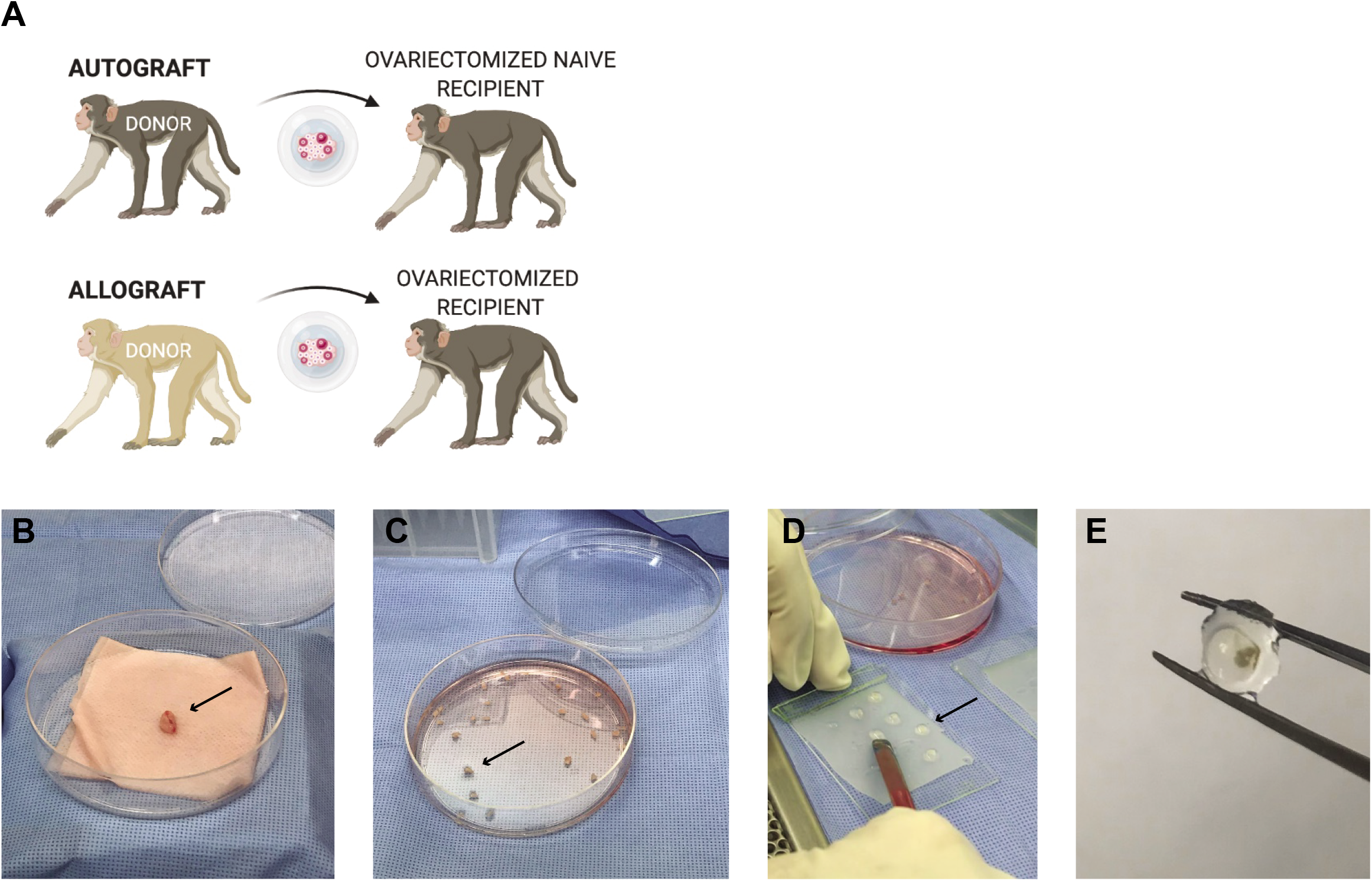
Harvest, processing, encapsulation, and explant of rhesus monkey ovarian tissue in immuno-isolating capsules. (**A**) Schematic for implantation of encapsulated auto- and allogeneic nonhuman primate ovarian tissue. (**B**) The ovary (arrow) was surgically removed from the donor animal and processed under sterile conditions. (**C**) The ovary was dissected into 1 mm3 (arrow) fragments and (**D**) encapsulated in immuno-isolating capsules (arrow). (**E**) Capsules were retrieved post-implantation after 4 or 5 months with no visible blood vessel infiltration or fibrosis surrounding the capsules. The ovarian tissue was visible in the center of the capsule.

### Follicles in the encapsulated ovarian autografts survive, resume folliculogenesis and result in detection of urinary estrone conjugate (E_1_C) and pregnanediol 3-glucuronide (PdG) in young rhesus monkeys

Consistent with the ovaries of peripubertal rhesus monkeys, the ovaries removed from the animals at the time of ovariectomy (Primates A and C) contained multiple primordial and primary follicles (**Figure 5A-D**). The cortical stroma had densely packed primordial follicles with oocytes surrounded by a single layer of flat squamous granulosa cells (**Figures 5Bi and 5Di**). In addition to primordial follicles, primary and a few small preantral follicles were also identified in the ovarian tissue. Primary follicles showed the characteristic oocyte surrounded by a single layer of cuboidal granulosa cells (**Figures 5Bii and 5Dii**), while multilayered secondary follicles (preantral follicle) had a few layers of granulosa cells surrounding the oocyte (**Figures 5Biii-iv and 5Diii-iv**). As anticipated in adolescent animals, antral follicles were not observed in the cortical fragments and the implanted fragments contained only primordial and primary follicles. After 4-5 months post-implantation the capsules with autologous tissue were removed, fixed, and analyzed for follicular development by histological analysis. All stages of follicles ranging from primordial to antral were observed in the capsules containing autografts (**Figure 5E-I**). Importantly, the presence of healthy-appearing and growing follicles supported continued folliculogenesis and is consistent with basal steroidogenesis.

**Figure 5:**
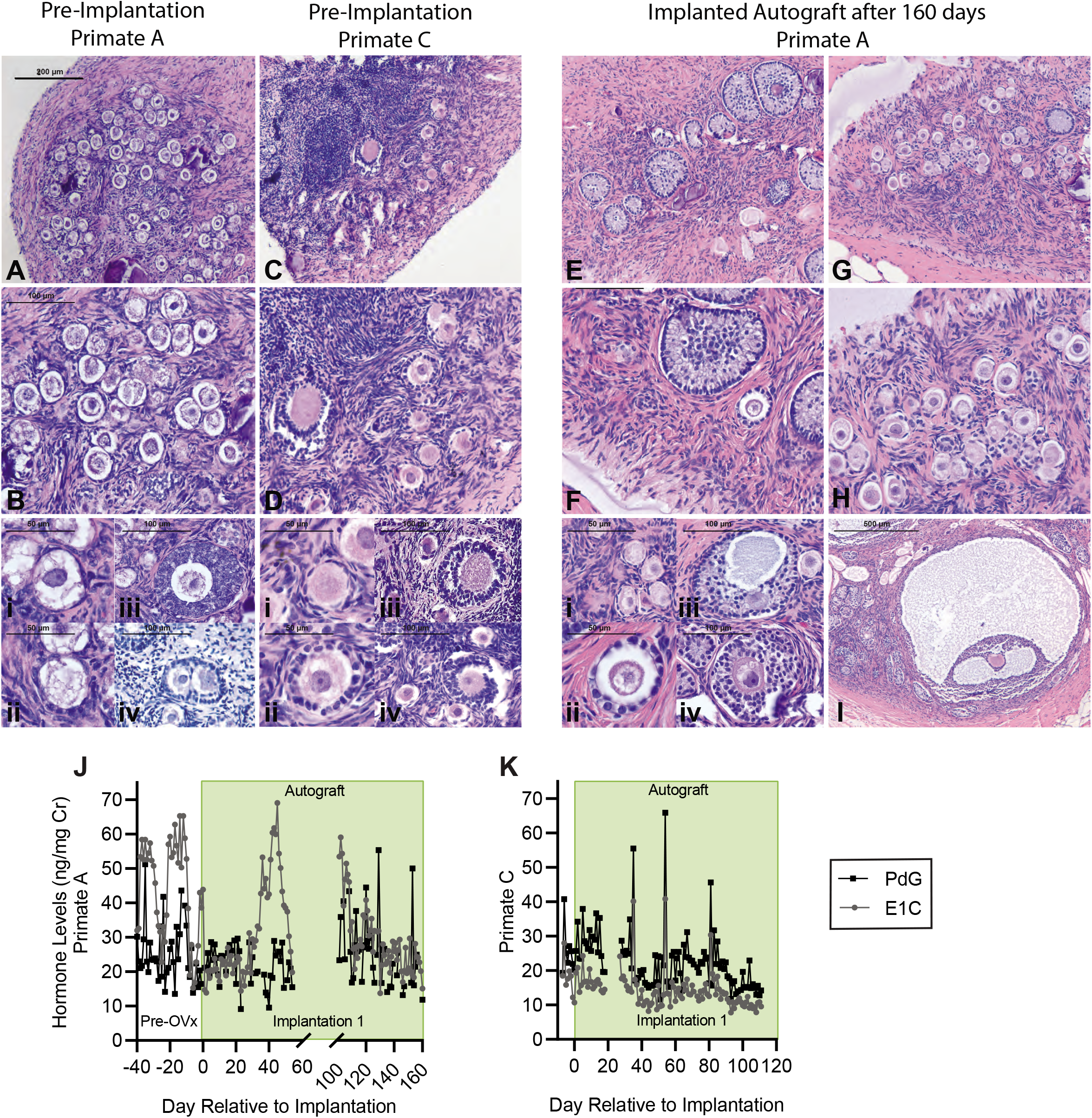
Histologic analysis of rhesus monkey donor ovarian tissue at implantation for Primate A (**A**, **B**, **Bi-iv**) and Primate C (**C**, **D**, **Di-iv**); autologous rhesus ovarian tissue encapsulated in immuno-isolating capsules for 5 months (**E**, **F**, **G**, **H**, **Hi-iv** and **I**). Scale bars: 200 μm (**A**, **C**, **E** and **G**), 100 μm (**B**, **D**, **F** and **H**), 50 μm (**i**, **ii**), 100 μm (**iii**, **iv**), 500 μm (**I**). Urinary estrone conjugate (E_1_C) and pregnanediol 3-glucuronide (PdG) levels in animals that received (**J**) encapsulated autologous tissue for 5 months, and (**K**) encapsulated autologous tissue for 4 months. Levels are normalized to urinary creatinine (Cr) ^48^.

We evaluated restoration of ovarian endocrine function by measuring urinary levels of E_1_C and PdG daily (no urine was collected in Primates A and B between days 55-100, and Primate C between days 20-27). E_1_C and PdG levels were normalized to urinary creatinine (Cr) levels as previously reported ^47,48^. Before ovariectomy all animals exhibited basal ovarian activity with levels of E_1_C and PdG fluctuating between 10-51 ng/mg Cr for E_1_C and 14-63 ng/mg Cr for PdG (**Figures 5J, K and 6B**, before Day 0). After ovariectomy followed by immediate implantation of autologous ovarian tissue encapsulated in immuno-isolating capsules, the levels of E_1_C and PdG were 10-30 ng/mg Cr. In Primate A (**Figure 5J**), the levels of E_1_C increased 25 days post-implantation reaching 50-70 ng/mg Cr. PdG also increased reaching 55 ng/mg Cr, similar to the measured levels of E_1_C and PdG before ovariectomy. Primate C with autologous implants showed lower levels of circulating E_1_C and PdG before ovariectomy (**Figure 5K**) ranging between 10-28 ng/mg Cr for E_1_C and 20-40 ng/mg Cr for PdG. After ovariectomy and immediate implantation of encapsulated ovarian autografts, the levels of E_1_C and PdG remained in the same range for 35 days, then fluctuated, reaching peak values at 40 ng/mg E_1_C and 65 ng/mg PdG. In both animals with encapsulated ovarian autografts the levels of E_1_C and PdG fluctuated from 6-69 ng/mg and 8-65 ng/mg Cr (**Figures 5J and K**) respectively, suggesting the encapsulated tissue was producing ovarian hormones at a basal level. A decline and peak approximately every 7 days was considered typical for fluctuations of basal steroid levels.

**Figure 6:**
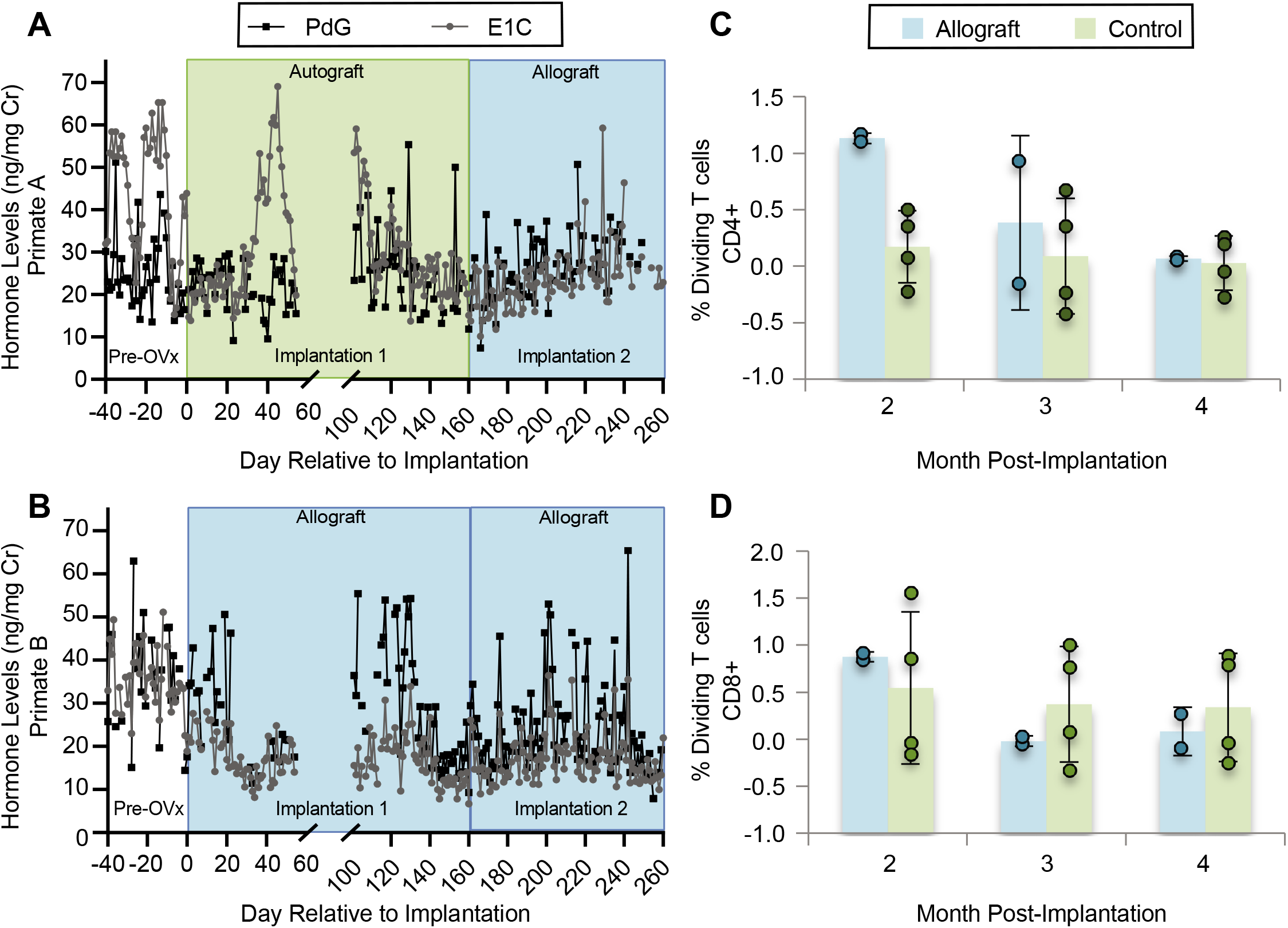
Urinary E1C and PdG levels and mixed lymphocyte culture in rhesus monkeys. Urinary E1C and PdG levels in rhesus monkeysthat received (**A**) encapsulated autologous tissue for 5 months followed by encapsulated allografts for 4 months, (**B**) encapsulated ovarian allografts for 5 months during the first implantation followed with a second implantation of encapsulated allografts for 4 months. Levels are normalized to urinary creatinine. Mixed lymphocyte culture of recipient cells reacted with donor cells quantifying dividing (**C**) CD4+ and (**D**) CD8+ T cells. Bar graphs represent mean±SD.

### Implanted encapsulated ovarian allografts secrete E_1_C and PdG and elicit minimal immune responses

Female rhesus monkeys implanted with encapsulated ovarian allografts exhibited basal ovarian activity with fluctuating levels of urinary E_1_C and PdG between 15 and 65 ng/mg Cr (E_1_C), and 13 to 63 ng/mg Cr (PdG) (**Figure 5J, 6A and B**, before Day 0).

Post-ovariectomy and immediate implantation of allogeneic ovarian tissue encapsulated in immuno-isolating capsules, the levels of E_1_C and PdG declined to 8-25 and 11-22 ng/mg Cr, respectively, in Primate B. (**Figure 6B**). E_1_C increased 130 and 200 days post-implantation reaching 34 and 36 ng/mg Cr in the first and second rounds of implantation respectively, with decline and peak approximately every 7 days. PdG also increased reaching 55 ng/mg Cr, similar to the measured levels of urinary E_1_C and PdG prior to ovariectomy. After 160 days, the encapsulated autografts from Primate A were explanted and encapsulated allografts were implanted (**Figure 6A**). The levels of circulating E_1_C and PdG initially downtrended to 10 and 28 ng/mg Cr for E_1_C and 7 to 37 ng/mL Cr for PdG. After implantation of encapsulated ovarian allografts, the levels of E_1_C and PdG continued to increase reaching 60 ng/mg Cr (E_1_C) at day 229 and 50 ng/mL Cr (PdG) at day 216 post-ovariectomy. Between the distinct peaks, the ovarian hormone levels appeared pulsatile but at a lower amplitude.

### Ovarian tissue encapsulated in PEG capsules did not elicit immune response

To assess whether the rhesus monkeys mounted an immune response against the encapsulated tissue, mixed lymphocyte culture assays were performed at the conclusion of the second round of implantations for all three animals. Peripheral blood mononuclear cells (PBMCs) from each animal were tested for reactivity to autologous cells (negative control) or to cells from the ovarian allograft donor. We observed no significant proliferation of CD4^+^ T cells in response to allogeneic donors when compared to proliferation in response to autologous or third-party cells (negative controls) (**Figure 6C**). Similarly, there was no significant spike in dividing CD8^+^ T cells in any of the animals when implanted with encapsulated allogeneic or autologous ovarian tissue. This finding was comparable to the observations after mixing responder cells with irrelevant, non-donor stimulators (**Figure 6D**).

## Discussion

We have demonstrated in prior publications that the PEG-based immuno-isolating capsules consisting of a degradable core with a non-degradable shell supports survival and development of syngeneic and allogeneic ovarian tissue, leading to restoration of ovarian endocrine function in immune competent murine hosts ^37,38^. While we have shown that encapsulated allografts survive in an immune competent mouse, the question remained as to whether encapsulation preserves tissue function in a previously sensitized host. It is crucial to answer this question in anticipation of clinical application of ovarian tissue allo-transplantation as patients will likely require multiple implants and/or implantations to maintain ovarian endocrine function over a prolonged time period, and are likely to be sensitized to donors. Here, we investigated whether encapsulating ovarian allografts in PEG-based capsules protects against rejection following multiple implantations in previously sensitized murine hosts and in young ovariectomized rhesus monkeys, while still promoting folliculogenesis.

First, we tested whether consecutive implantations of non-encapsulated allogeneic ovarian tissue resulted in sensitization of the host, measured by allo-specific antibodies in the recipient’s serum. Hosts were sensitized by implantation of non-encapsulated ovarian allografts and the sensitization was confirmed by accelerated rejection of the second series of non-encapsulated allografts. The allo-specific IgG spiked 7 days after the second implantation, one week earlier than following the first implantation. Rejection was confirmed by tissue destruction and CD8^+^ T cell infiltrates. In contrast, we found that encapsulation of ovarian allografts avoids rejection in recipients sensitized against the donor. Thus, encapsulated grafts implanted in sensitized murine recipients restored ovarian function for up to 120 days after implantation, and the tissue was viable with allo-antibody levels at a plateau suggesting the sensitized mice did not respond further to allo-antigens.

As a next step, we investigated the ability of an immuno-isolating capsule to support follicular development and restore basal ovarian endocrine function in young ovariectomized rhesus monkeys implanted with autologous or allogeneic ovarian tissue. While we have previously shown that the capsule supported murine follicular development and protected ovarian allografts from rejection, the question remained whether the capsule could support the volumetric expansion of nonhuman primate follicles without compromising the capsule’s immuno-isolating capability. Importantly, to mimic the clinical scenario of puberty induction in young cancer survivors, the three animals in this study were young, peripubertal females that did not yet exhibit regular follicular and luteal phases of the menstrual cycle. Once the monkeys were ovariectomized and implanted with encapsulated ovarian tissue, they maintained basal levels of E_1_C and PdG throughout the entirety of the implantation period similar to the hormone levels detected prior to ovariectomy. While there was no indication of ovarian cycles expected in mature females, the presence of basal ovarian hormones following the implantations reflected follicular activation, follicle recruitment into the growing pool, and progression to the small antral stage, particularly in the autograft implant ^49^.

To confirm whether the PEG capsules could support nonhuman primate folliculogenesis, histological analysis was performed. Healthy follicles were consistently present in the encapsulated autologous ovarian implants. We observed the presence of all follicular stages (up to the antral stage), indicating folliculogenesis was supported in the capsules. The presence of small antral follicles suggest that nonhuman primate ovarian tissue can undergo the necessary volumetric expansion during folliculogenesis to the small antral stage without capsule restrictions, most likely due to the degradation of the core and the viscoelastic properties of the shell. Upon visual evaluation after explant, there was no evidence of fibrous tissue or inflammation at the site, similar to our prior findings in rodents ^35^.

To determine whether implanted young rhesus monkeys mounted an immune response against the encapsulated allografts, mixed lymphocyte cultures were performed. We observed that when PBMCs from the animals were exposed to cells from the donor, there was no significant CD4^+^ and CD8^+^ T cell proliferation when compared to autologous or third party controls. This finding suggests that the animals were not sensitized to the allo-antigens and the capsules isolated the ovarian allografts from the immune system of the recipient. Importantly, the PEG capsules were able to support folliculogenesis without compromising the immuno-isolating capability.

We believe this report is the first to demonstrate the use of an immuno-isolating capsule to support nonhuman primate folliculogenesis in a non-vascularized ovarian tissue graft, while protecting the ovarian allograft from the immune system, and maintaining basal levels of ovarian endocrine function. Whether restoration of cyclic ovarian function typical of adult female rhesus monkeys during spontaneous menstrual cycles can be supported by the immuno-isolating capsules remains to be determined.

In summary, we have established that encapsulating allogeneic ovarian tissue in a PEG-based capsule protects the allograft and restores and maintains some ovarian endocrine function, prevents the sensitization of the host immune system, and functions similarly in sensitized and naïve mice as well as rhesus monkeys. Further studies assessing the ability of the dual-layered PEG capsule to support normal ovarian cyclicity, including corpus luteum formation, will be important to further verify that the immuno-isolation construct is functional. Further capsule modifications to support larger graft fragments may also be required to promote vasculature formation around the capsule and greater diffusion of essential metabolites and nutrients without the risk of immune rejection.

In patients experiencing POI, allogeneic cell-based therapy has the potential to restore hormonal function in a physiologic pulsatile and self-regulating manner, potentially avoiding side effects of pharmacological HRT as well the risk of cancer recurrence associated with auto-transplantation.

Additional populations that could benefit from encapsulated ovarian allo- or auto-transplantation include women with genetic causes of POI or women with autoimmune diseases such as lupus that receive cyclophosphamide immunosuppressive therapy. The ability of the immuno-isolating dual-layered PEG capsule to avoid sensitization and sustain prolonged function in sensitized hosts is paramount to allowing sequential grafts since children or young adults with POI will require treatment that may span decades.

## Materials and Methods

### Animals

All animal procedures conformed to the requirements of the Animal Welfare Act and protocols were approved prior to implementation by the Institutional Animal Care and Use Committee (IACUC) at the University of Michigan for the rodents (PRO00007716) and University of California, Davis for rhesus monkeys (see below).

Activities related to nonhuman primate animal care (diet, housing) were performed per IACUC approved California National Primate Research Center standard operating procedures (SOPs). Physical signs were monitored daily, and body weights were assessed each time the animals were sedated with ketamine (10 mg/kg).

### Experimental Design

#### Murine model

Adult female mice (BALB/c) underwent bilateral ovariectomies to induce POI. As a positive control for sensitization, BALB/c mice were implanted subcutaneously with a non-encapsulated ovary from 6-8 day-old CBA × C57BL/6 (F1) mice for 42 days followed by a subsequent implantation of another non-encapsulated F1 ovary for 14 days (**Figure 1A)** (n=7 mice). For experimental groups, BALB/c mice were (i) implanted with encapsulated ovarian allogeneic (F1) tissue for 60 days followed by a subsequent implantation of encapsulated ovarian allografts for another 60 days (n=5 mice) or (ii) implanted with non-encapsulated F1 ovarian tissue for 60 days to induce sensitization followed by a subsequent implantation of encapsulated ovarian allografts for 60 days (n=3 mice). A total of 15 mice were used in these studies. Each mouse was implanted with two replicate grafts or non-encapsulated grafts. Bilateral ovariectomies were performed on adult female mice (BALB/c) at 12-16 weeks of age. The mice were anesthetized by isoflurane for the surgical procedure and Carprofen (5 mg/kg body weight, Rimadyl, Zoetis) was administered subcutaneously for analgesia. The intraperitoneal space was exposed through a midline incision in the abdominal wall secured using an abdominal retractor. The ovaries were removed, and the muscle and skin layer of the abdominal wall were closed with 5/0 absorbable sutures (AD Surgical). The mice recovered in a clean warmed cage and received another dose of Carprofen 12 hours post recovery or as needed.

### Collection of Donor Murine Ovaries

Ovaries from 6 to 8 day-old CBA × C57BL/6 (F1) mice were collected and transferred to Leibovitz L-15 media (P/N 11415, Gibco, USA). The ovaries were dissected into 2–4 pieces and transferred into maintenance media (α-MEM, P/N 32561, Gibco, USA) and placed in a CO_2_ incubator prior to encapsulation.

### Hydrogel Preparation and Murine Ovarian Tissue Encapsulation

The degradable core of the PEG-based capsule was prepared with 8-arm PEG-VinylSulfone (PEG-VS) (40kDa, Jenkem Technology, Beijing, China) and cross-linked with a plasmin sensitive tri-functional peptide sequence (Ac-GCYK↓NSGCYK↓NSCG, MW 1525.69 g/mol, >90% Purity, Genscript, ↓ indicates the cleavage site of the peptide). The non-degradable shell of the capsule was prepared using 4-arm PEG-VS (20kDa, Jenkem Technology, Beijing, China) with Irgacure 2959 (BASF, Switzerland, MW=224.3) and 0.1% N-vinyl-2-pyrrolidone (P/N V3409, Sigma-Aldrich, St. Louis, USA). The detailed protocol is described in Day et al. ^33,37^. The murine ovarian fragments were first encapsulated in 4 μL of the degradable PEG hydrogel for 5 minutes. The degradable core and graft were then placed in the center of a 10 μL droplet of PEG-VS precursor solution (5% w/v PEG-VS, 0.4% Irgacure 2959, 0.1% NVP) and exposed to UV light for 6 minutes. All constructs were imaged immediately after encapsulation of the tissue to confirm encapsulation.

### Subcutaneous Implantation in Mice

A small incision was made on the dorsal side of anesthetized mice (BALB/c) and the immuno-isolating capsules with the ovarian allografts or non-encapsulated ovarian allografts were implanted subcutaneously. The skin was closed using 5/0 absorbable sutures. The mice received Carprofen for analgesia for at least 24 hours after surgery or as needed.

### Vaginal Cytology in Mice

Daily vaginal cytology was performed after ovariectomy to assess estrous cycle status and confirm cessation for seven days post-surgery. Starting one week after allograft implantation, vaginal cytology was performed daily to assess estrus cycle status and determine if estrous cyclicity resumed until euthanasia. Observation of the transition from leukocytes to cornified cells at least once a week was the criteria used to determine a resumed or continued cycle.

### Flow Cytometry of Mouse Serum

Serum allo-antibody titer measurements were performed using flow cytometry before and after implantation and reported as MFI for the highest dilution showing fluorescence detectable above background (non-immune serum from a non-implanted mouse) in immunized mice (positive controls implanted with non-encapsulated ovary allografts). Thymocytes were isolated from CBA × C57BL/6 donor mice and incubated with serially diluted recipient serum for 30 minutes at 4°C. Antibodies bound to the thymocytes were detected by Cy5-conjugated goat anti-mouse IgG (1:250 dilution, 1030-15, Southern Biotech, Birmingham, AL) and Alexa Fluor 488-conjugated goat anti-mouse IgM (1:250 dilution, 1020-30, Southern Biotech, Birmingham, AL) for 30 minutes at 4°C and analyzed in a BD FACSCanto II (BD Biosciences, Franklin Lakes, NJ). The MFI in the APC-channel (measuring bound IgG) and FITC channel (measuring bound IgM) were determined with FlowJo 10 software (FlowJo, LLC, Ashland, OR).

### Histological Analysis of the Murine Ovarian Allografts and Encapsulated Ovarian Allografts

Following euthanasia, the immuno-isolating capsules or allografts were retrieved from the animals, fixed in 4% paraformaldehyde at 4°C overnight, then transferred and stored in 70% ethanol at 4°C. Samples were histologically processed for paraffin embedding, serially sectioned at 5 μm thickness, and stained with hematoxylin and eosin (H&E).

### Immunohistochemistry (IHC) of Mouse Ovarian Allografts

To analyze T cell infiltration into non-encapsulated or encapsulated ovarian allografts, paraffin-sections were stained to identify CD4^+^ and CD8^+^ T cells. Following deparaffinization with xylene and rehydration, the sections were incubated in antigen retrieval buffer, pH 9.0 (ab94681, Abcam, Cambridge, MA) for 20 minutes at 97°C and an additional 20 minutes at room temperature to cool. Next, the slides were incubated with KPL Universal Block (5560-0009, SeraCare Life Sciences, Milford, MA) to block non-specific binding sites for 30 minutes at room temperature. The sections were incubated at room temperature for 1 hour with primary antibodies: rabbit monoclonal anti-mouse CD4 antibody (1:1000 dilution, ab183685, Abcam, Cambridge, MA), and rabbit polyclonal anti-mouse CD8 antibody (1:500 dilution, ab203035, Abcam, Cambridge, MA). The slides were subsequently incubated at room temperature with secondary antibodies: goat anti-rabbit Ig (1:100 dilution for 15 minutes, 4010-05, Southern Biotech, Birmingham, AL) for CD4 and goat anti-rabbit (1:50 dilution for 30 minutes, 4010-05, Southern Biotech, Birmingham, AL) for CD8. Diaminobenzidine (DAB) (BDB2004L, Betazoid DAB Chromogen kit, BioCare Medical, Pacheco, CA) was used as a chromogen for 10 minutes at room temperature. Hematoxylin (220-102, Fischer Scientific) was used as a counterstain. For negative controls, paraffin sections were incubated without the primary antibody. To assess presence or absence of CD8^+^ T cells, 12 sections from the front, middle, and end of each specimen were examined, to represent the full thickness of the implant. Five equal sized fields in the 4 corners and center of each section were assessed for positive DAB chromogen staining.

#### Rhesus monkey model

### Ovariectomies and Subcutaneous Implantation in Recipient Rhesus Monkeys

Rhesus monkeys (~3 years of age, ~4 kg) were sedated with ketamine (10 mg/kg; IM) and prepared for bilateral ovariectomy according to established protocols^47^. Briefly, atropine was given (0.04 mg/kg) followed by intubation for the administration of isoflurane (to effect), and an indwelling catheter placed for intravenous (IV) fluids. A small midline incision was made, and the ovaries were exposed, removed individually, and placed in a sterile culture dish for processing in a biosafety cabinet under aseptic conditions. Ovarian tissue from Primate A was encapsulated in the immuno-isolating hydrogel under aseptic conditions and implanted in both Primate A (autograft) and Primate B (allograft). Once the encapsulated grafts were ready for implantation, a small incision (~0.5 cm) was made between the scapulae. The encapsulated tissue was gently inserted under aseptic conditions and the incisions were sutured closed and reinforced with skin glue. Blood and urine samples were collected regularly as described below. All animals were monitored post-surgery per SOP and administered analgesics post-operatively.

At approximately five months post initial implantation surgery, animals were prepared for a second implantation surgery. Primate C was ovariectomized and received fresh encapsulated autologous tissue grafts (right ovary) as described above. The first implantation round of encapsulated ovarian grafts were explanted from Primates A and B, which then received encapsulated allogeneic tissue from Primate C. At approximately four months post-implantation the encapsulated tissue was collected for analysis from all three animals.

Thus overall, two rounds of subcutaneous implantations of encapsulated autologous and allogeneic ovarian tissue fragments were performed. In the first round, one animal received 20 encapsulated autologous ovarian fragments and one animal received 20 encapsulated allogeneic ovarian fragments. In the second round three animals (two animals from the first round and an additional animal C) received 20 encapsulated ovarian fragments (N=1 autologous and N=2 allogeneic) (**Figure 4**).

### Primate Ovarian Tissue Encapsulation

The primate ovarian allograft tissue was encapsulated using the same capsule formulation as described above for the murine model. The primate ovaries were dissected into approximately 60 fragments measuring 1×1×1mm (1 mm^3^) and transferred into maintenance media (α- MEM, P/N 32561, Gibco, USA) prior to encapsulation (**Figure 4**). The monkey ovarian tissue fragments were first individually encapsulated in an 8 μL degradable PEG core, allowed to cross-link for 5 minutes, and were then placed in the center of a 20 μL droplet of PEG-VS precursor solution (5% w/v PEG-VS, 0.4% Irgacure 2959, 0.1% NVP) and exposed to UV light for 6 minutes.

### Primate Urinary E_1_C and PdG analysis

Urine samples were collected daily from cage pans according to SOP starting one month prior to surgery and through most of the post-implantation period ^47,48^. Rhesus monkeys were housed to ensure urine was collected from individual animals. Collected samples were centrifuged at 1,500 rpm for 10 minutes and supernatant (~2-3 ml) was transferred to a cryogenic vial (Caplugs Evergreen, Buffalo, NY) and stored at ≤ −20°C until processed. Urinary E_1_C and PdG levels were determined by the Endocrine Core (University of California – Davis) via the analysis of the urine supernatant according to established protocols ^48^.

### Mixed Primate Lymphocyte Culture

PBMCs were isolated from blood samples collected monthly by centrifugation onto a step gradient of Lymphocyte Separation Medium (LSM; MP Biomedicals, LLC, Solon, OH). Stimulator cells were treated with 40 μg/ml mitomycin C (Sigma) at 37°C for 30 minutes. CFSE-labeled responder cells were stimulated with an equal number of unlabeled stimulator cells or with plate-bound anti-CD3 antibody (positive control; clone SP34-2). Cells were harvested after 4-6 days of incubation, stained with fluorescently labeled antibodies including those specific for CD4 and CD8 (clones L200 and 3B5, respectively); cytometry data were collected on a BD Fortessa; and the results were analyzed in FlowJo’s proliferation platform (FlowJo, LLC, Ashland, OR).

### Histological Analysis of the Encapsulated Ovarian Primate Allografts

Following euthanasia, the immuno-isolating capsules with ovarian tissue grafts were retrieved from the animals, fixed in 4% paraformaldehyde at 4°C overnight, transferred and stored in 70% ethanol at 4°C. During the encapsulation process of the ovarian grafts, fresh non-encapsulated ovarian tissue samples from each donor were also fixed and stored similarly. Samples were histologically processed for paraffin embedding, serially sectioned at 5 μm thickness, and stained with H&E.

### Statistics

Statistical analysis was performed using GraphPad Prism software version 9.0.0. Multiple comparisons were made using One-way ANOVA. Significance was determined by Tukey’s multiple comparisons test.

## Acknowledgements

Figure schematics in Figures 1a, 1d, 2a, 2b, 3a, 4 were created with BioRender.com.

The authors would like to acknowledge the assistance and expertise of B. Lasley (core lead, Endocrine Core, University of California – Davis) in the analysis of urinary E_1_C and PdG.

## Author Contributions

AS, MC and AT conceived the study idea. AS, MC, AT, JRD, AD developed the methodology, executed the experiments and collected samples. JRD, CLF, AD, DJHO, MGMB, MLM, CL, JB, EF, MZ processed the samples, performed the formal analysis and data visualization. AS, MC, AT were responsible for project administration, supervision and funding acquisition. JRD, AD, AS and MC prepared the original draft of the manuscript. All authors reviewed and edited the manuscript.

